# Seeking an optimal dosing regimen for OZ439-DSM265 combination therapy for treating uncomplicated falciparum malaria

**DOI:** 10.1101/2020.11.02.366112

**Authors:** Saber Dini, Sophie G Zaloumis, David J Price, Nathalie Gobeau, Anne Kümmel, Mohammed Cherkaoui, Joerg J Moehrle, James S McCarthy, Julie A Simpson

## Abstract

The efficacy of Artemisinin-based Combination Therapies (ACTs), the first-line treatments of uncomplicated falciparum malaria, has been declining in malaria endemic countries due to the emergence of malaria parasites resistant to these compounds. Novel alternative therapies are needed urgently to prevent the likely surge in morbidity and mortality due to failing ACTs. This study investigates the efficacy of the combination of two novel drugs, OZ439 and DSM265, using a biologically informed within-host mathematical model that accounts for the pharmacodynamic interaction between the two drugs. Model parameters were estimated using data from healthy volunteers infected with falciparum malaria collected from four trials: three that administered OZ439 and DSM265 alone, and the fourth a combination of OZ439-DSM265. Posterior predictive simulations of the model were performed to determine efficacious dosing regimens. One such regimen that predicted at least 90% of infected individuals cured 42 days after the administration of the drugs, while within the tolerable dose range, is 800 mg of OZ439 and 450 mg of DSM265. Our model can be used to inform future phase 2 and 3 clinical trials of OZ439-DSM265, fast-tracking the deployment of this combination therapy in the regions where ACTs are failing.

## Introduction

Artemisinin-based Combination Therapies (ACTs) have been the first-line treatment of uncomplicated falciparum malaria in most malaria-endemic countries for more than two decades [1]. During this period, ACTs have played a central role in malaria control and the decline in clinical cases and malaria attributable deaths. Alarmingly, the efficacy of ACTs has declined below 50% in some regions [2], due to the emergence and spread of parasites resistant to the artemisinins across the Greater Mekong Region [3, 4, 5]. This worrying trend threatens to reverse the recent progress against *Plasmodium falciparum* (*P. falciparum*) achieved by widespread availability of ACTs and, of greater concern, highlights the prospect of untreatable falciparum malaria in the absence of efficacious alternative antimalarial treatments.

Various alternative treatments have been suggested, such as combining a failing ACT with an already available partner drug, known as Triple Artemisinin-based Combination Therapy (TACT) [6, 2], or producing novel synthetic antimalarials [7]. Key features of a successful treatment include a dosing regimen that is highly effective and easy to adhere to, so that sub-therapeutic concentrations are avoided, and combining drugs with different modes of action to prevent the development of resistance to each individual drug [8, 9].

OZ439 (also known as artefenomel) is a novel antimalarial drug with a mechanism of action similar to artesunate, i.e. activation of an endoperoxide bond which in turn damages various proteins of the parasite using free radicals and reactive intermediates [10]. However, unlike artesunate which has an elimination half-life of *∼*1 h (500 mg dose) [11], OZ439 features a significantly longer elimination half-life (*∼*260 h (500 mg dose) [12]), thus exposing the parasites to OZ439 concentrations for a long duration after a single dose. The favourable pharmacokinetic (PK) properties of OZ439 as well as its safety and tolerability at relatively high doses, and *in vitro* data suggesting that it is active against artemisinin-resistant parasites [13] make it a potential candidate to replace the artemisinins in regions where resistance to this drug has risen.

DSM265 is another novel synthetic antimalarial drug with a long elimination half-life (between 86 and 118 h) and satisfactory safety and tolerability [14, 15]. Similar to OZ439, the long presence of this drug in the blood plasma allows administration of a single-dose regimen, whereas current dosing regimens for ACTs recommend daily administration (and for artemether-lumefantrine twice daily) for three days. DSM265 kills the parasites by inhibiting *Plasmodium* dihydroorotate dehydrogenase (DHODH) which is a vital enzyme for pyrimidine biosynthesis of the parasite [16]. None of the currently administered antimalarials has this mechanism of action, making DSM265 an attractive candidate for a new antimalarial drug.

The promising pharmacological characteristics of OZ439 and DSM265 described above suggest that these drugs may be suitable candidates as a combination antimalarial treatment. A recent trial evaluating the OZ439-DSM265 combination in healthy volunteers infected with blood-stage falciparum malaria found satisfactory safety and tolerability and promising antimalarial activity [7]. Relatively low doses were intentionally administered in the trial to allow parasitaemia recrudescence which provides important insights into the parasitological responses and is more informative for pharmacometric studies. Subsequent trials are required to investigate the efficacy of higher doses of OZ439 and DSM265 in this combination treatment. A selected efficacious dosing regimen must also satisfy safety and tolerability constraints – both drugs have shown good safety and tolerability profiles up to relatively high does [15, 17]. In addition, the exposure profiles of drugs must overlap to a large extent to reduce the likelihood of resistance selection by the parasites due to their exposure to sub-therapeutic levels of only one drug.

This study focuses on the efficacy of the OZ439-DSM265 compound using a biologically informed pharmacodynamic (PD) mathematical model that accounts for the stage-specific killing action of the drugs [18, 19, 20, 21]. The PD interaction between OZ439 and DSM265 was determined and accommodated in the model. Data from four separate trials of healthy volunteers inoculated with blood-stage falciparum malaria (OZ439 and DSM265 given alone and the OZ439-DSM265 combination therapy) were analysed. A Bayesian approach was used for parameter estimation which enabled us to use the information from modelling the mono-therapy trials data as prior information for estimating the parameters of the combination therapy. Simulations of the fitted model at different OZ439-DSM265 doses were performed to propose regimens required to cure (within 42 days of follow-up) at least 90% of individuals infected with uncomplicated *P. falciparum*. This work was aimed at informing the selection of dose regimens to investigate in future clinical trials of the OZ439-DSM265 combination.

## Results

Data collected from volunteers infected with *P. falciparum* in mono-therapy trials of OZ439 and DSM265 and the OZ439-DSM265 combination therapy trial were used for model fitting; Table 1. The measured drug concentrations of the volunteers were first used to estimate the population PK model parameters and between-subject variability. A sequential PK-PD modelling approach was then performed where PK profiles were simulated for each volunteer based on the post-hoc individual PK parameter estimates, which were subsequently substituted into the PD model; see the *Materials and Methods* section for further details.

**Table 1:** Dosing regimens of OZ439 and DSM265 administered in four clinical trials of volunteers infected with *P. falciparum*.

| Treatment* | Cohort | Dose [mg] | Drug administration time* [hours] | Parasitaemia [no. parasites/mL] at time 0 median (range) |
| --- | --- | --- | --- | --- |
| OZ439 [12] | A (n=8) | 100 | 0 | 5487 (939–7517) |
|  | B (n=8) | 200 | 0 | 5676 (407–24152) |
|  | C (n=8) | 500 | 0 | 4475 (953–10315) |
| DSM265† [14, 15] | A (n=7) | 150 | 0 | 8442 (1676–13286) |
|  | B (n=7) | 400 | 0 | 8457 (1235–62044) |
| OZ439-DSM265 [7] | A (n=8) | OZ439 200 | 0 | 1629 (289–7678) |
|  |  | DSM265 100 | 2 |  |
|  | B (n=5) | OZ439 200 | 0 | 12364 (3495–27312) |
|  |  | DSM265 50 | 2 |  |

\* See the *Data* section for further information about inoculation of the volunteers with *P. falciparum* and their treatment.
† Data from two separate trials (Cohorts A [15] and B [14]) for DSM265 were used.

### Pharmacokinetics

A two-compartment PK model with first and zero order absorption rates best described the OZ439 and DSM265 drug concentrations, respectively (*Materials and Methods* section). Table 2 summarises the estimated PK parameters for the data from the combination therapy trial. Figure 1 shows the observed and simulated drug concentration profiles of the 13 volunteers; see Figure S1 of Supplementary Material for the PK profiles of the volunteers receiving OZ439 and DSM265 alone. The profiles show significantly higher concentration of DSM265 in the blood plasma of volunteers compared with OZ439.

**Table 2:** Parameter values of the PK models. A two compartment model with zero-order and first-order absorption were used for DSM265 and OZ439, respectively.

| Parameter | Drug | Estimate (RSE) | BSV (RSE) | Description |
| --- | --- | --- | --- | --- |
| $F$ | DSM265 | 1 (Fixed) | 0.0897 (44.50%) | Relative bioavailability |
|  | OZ439 | 1 (Fixed) | 0 (Fixed) |  |
| $CL/F$ | DSM265 | 0.514 (3.20%) | 0.232 (11%) | Apparent clearance [L/hour] |
|  | OZ439 | 43.2 (9.60%) | 0.261 (16.40%) |  |
| $V_c/F$ | DSM265 | 12.4 (14%) | 0.66 (17.20%) | Apparent central volume [L] |
|  | OZ439 | 148 (12.50%) | 0 (Fixed) |  |
| $Q/F$ | DSM265 | 38.5 (8.24%) | 0.25 (Fixed) | Apparent intercompartmental clearance [L/hour] |
|  | OZ439 | 11.1 (8.66%) | 0 (Fixed) |  |
| $V_p/F$ | DSM265 | 61.8 (4.29%) | 0.25 (Fixed) | Apparent peripheral volume [L] |
|  | OZ439 | 903 (12.20%) | 0.276 (38.80%) |  |
| $T_{k0}$ | DSM265 | 2.64 (5.86%) | 0.413 (10.20%) | Absorption time [hours] |
| $k_a$ | OZ439 | 0.198 (6.85%) | 0.106 (58.60%) | Absorption rate parameter [1/hour] |
| $T_{lag}$ | DSM265 | 0 (Fixed) | 0 (Fixed) | Absorption lag time [hours] |
|  | OZ439 | 0.418 (2.92%) | 0.107 (20.70%) |  |
| $\beta_F$ | DSM265 | -0.114 (22.20%) | - | Coefficient of dose on $F$<br>Reference dose: 400mg |
| $\beta_{V_p}$ | DSM265 | 1.61 (16.90%) | - | Coefficient of weight on $F$<br>Reference weight: 76.8kg |
| $\beta_{CL}$ | DSM265 | 0.649 (34.10%) | - | Coefficient of dose on $CL$<br>Reference dose: 500mg (DSM265)<br>400mg (OZ439) |
|  | OZ439 | -0.473 (18.20%) | - |  |

BSV: between subject variability
RSE: relative standard error defined as (standard deviation/mean) × 100

**Figure 1:**
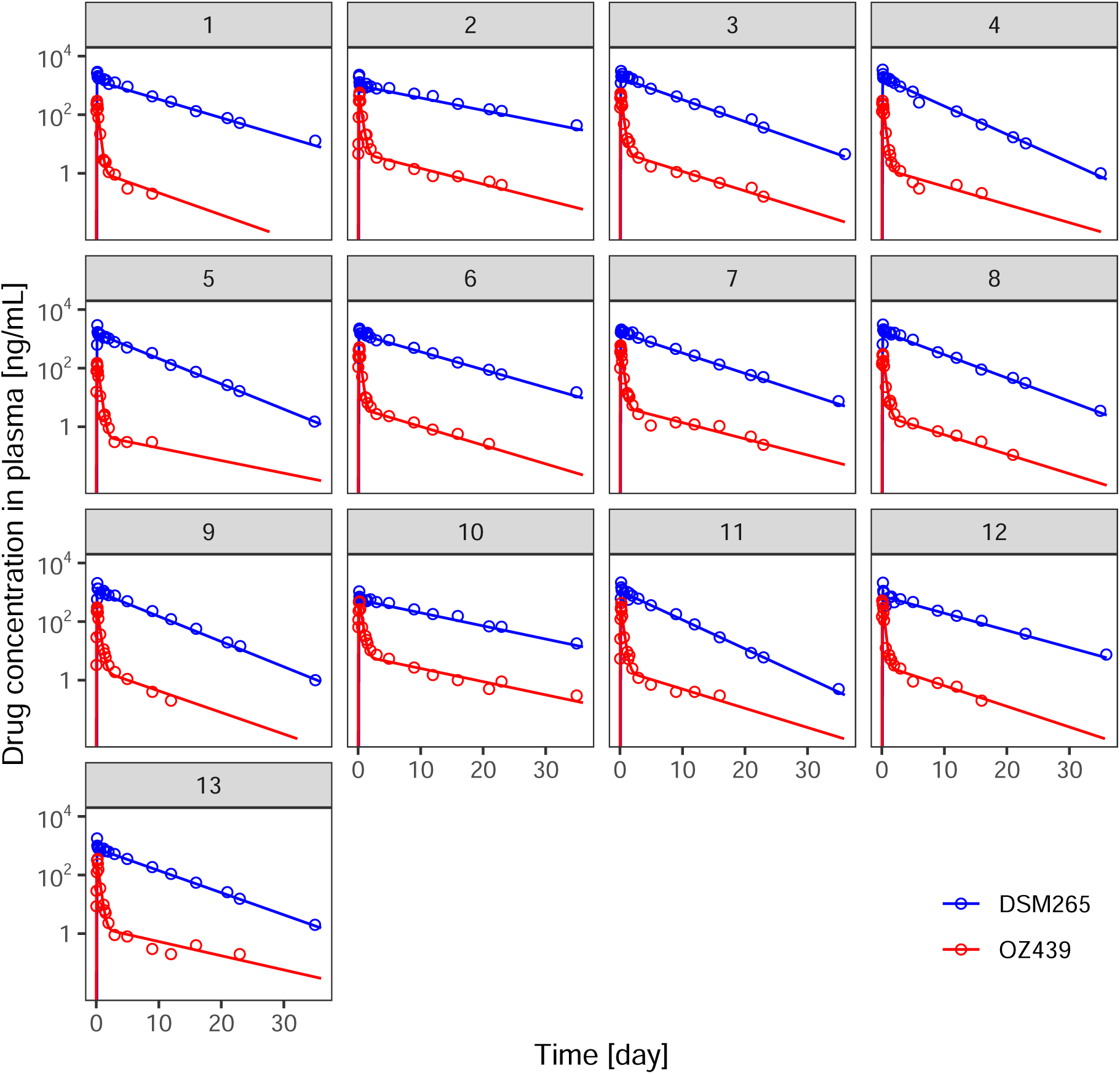
Pharmacokinetic profiles of OZ439 and DSM265 for the OZ439-DSM265 combination therapy. The points show the measured drug concentrations and the lines are the generated simulations using the mode of the conditional distribution of the individual PK parameters – the population parameters are listed in Table 2. A two-compartment model with first and zero order absorption rates were used for OZ439 and DSM265, respectively. Volunteers 1–8 received 200 mg of OZ439 and 100 mg of DSM265, and volunteers 9–13 received 200 mg of OZ439 and 50 mg of DSM265.

### Pharmacodynamics

The antimalarial activities of the drugs were modelled using the mathematical model defined in equation (1). The model accounts for differential action of the drugs on different stages of the parasite life cycle and PD interaction between the drugs. Initially, the PD model was fitted to the measured parasitaemia of volunteers in the OZ439 and DSM265 mono-therapy trials (results shown in Section *Mono-therapies* in the Supplementary Material). The estimated posterior distributions were then used to inform the estimation of *E*_*max*_, *γ, EC*_50_ and *k*_*e*0_ in the OZ439-DSM265 model fitted to the data of the combination therapy trial (see the *Materials and Methods* section).

Figure 2 shows the observed parasitaemia profiles for the 13 volunteers (black lines and dots), overlaid with the posterior predictive distributions (red line: median; shaded region: 95% credible interval). The results show that the PK-PD model captures the dynamics of the observed parasitaemia well for all individuals.

**Figure 2:**
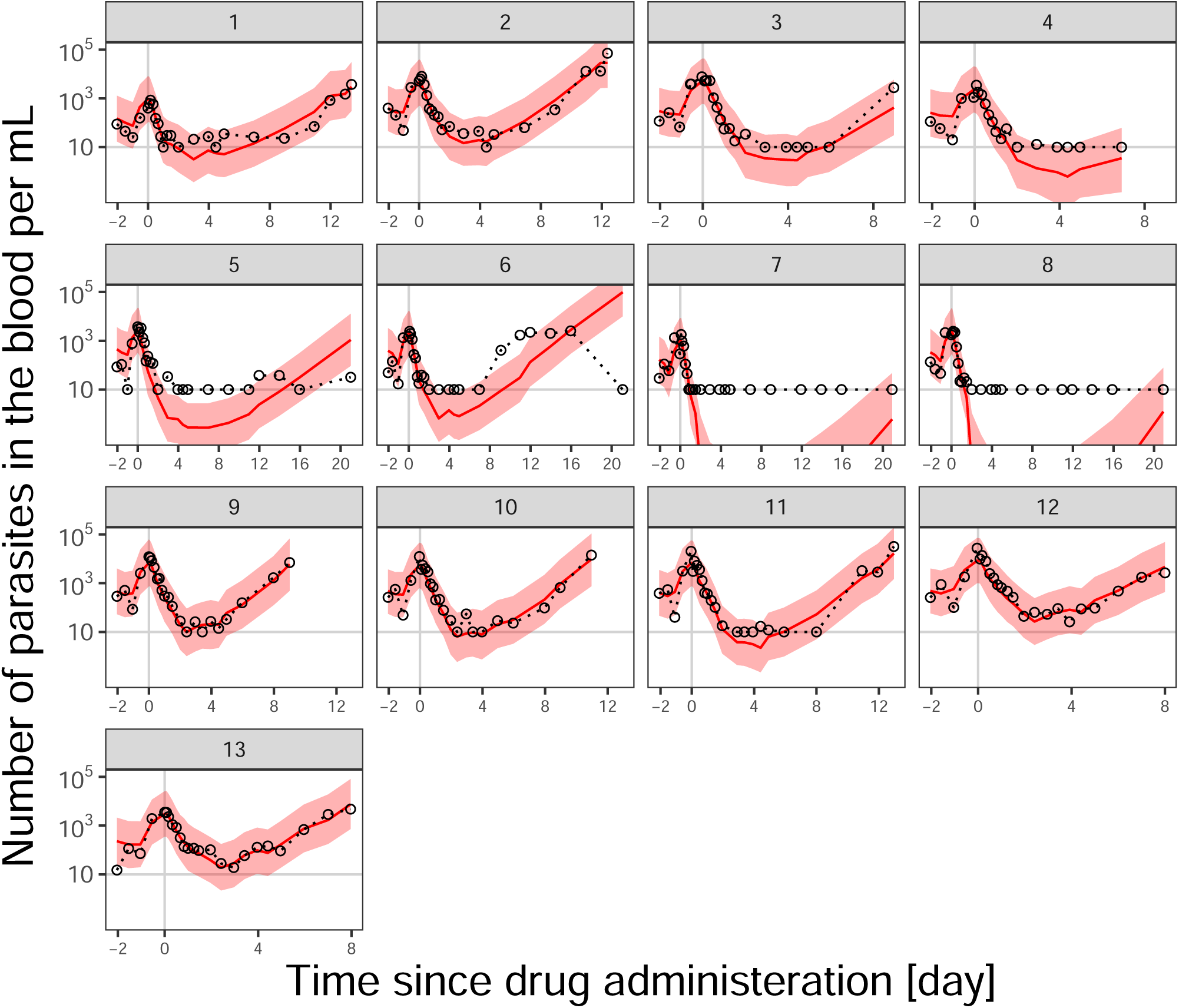
Posterior predictive check of the within-host PK-PD model fitted to the parastaemia of volunteers in the combination therapy trial. The black circles are the observed parasitaemia; the red line and the shaded area denote the median and 95% credible interval (between 2.5^th^ and 97.5^th^ percentiles) of 8,000 simulations of the model using the posterior samples of the individual PD parameters (see *Materials and Methods* section). Volunteers 1 to 8 belong to Cohort A (OZ439: 200 mg; DSM265: 100 mg) and volunteers 9 to 13 to Cohort B (OZ439: 200 mg; DSM265: 50 mg); see Table 1 for details of the cohorts. The grey vertical line at time = 0 shows the administration time of OZ439; DSM265 was administered 2 hours after the OZ439 administration; see Table 1 The Lower Limit of Quantification (LLOQ), shown with the horizontal line, was 10 [parasites/mL].

The estimated PD parameters for the combination therapy are summarised in Table 3; Tables S1 and S2 of the Supplementary Material include the details of the posterior distributions of the PD parameters estimated from modelling of the mono-therapy trials data. 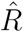 and *n*_*eff*_ metrics (see Table S3 of the Supplementary Material) indicate the convergence of the Hamiltonian Monte Carlo (HMC) Markov chains: 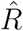 values are close to one, and *n*_*eff*_ values are large; see *Materials and Methods* section. Other model fit diagnostics for the four trials are illustrated in Section 4 of the Supplementary Material.

**Table 3:** Estimated parameter values of the PD model fitted to the data of OZ439-DSM265 combination therapy.

| Parameter* | Drug | Prior bounds<sup>†</sup> | Posterior median<sup>‡</sup><br>(95% credible interval) | BSV<sup>§</sup><br>(95% credible interval) |
| --- | --- | --- | --- | --- |
| $N_0$ | | $(5.04 \times 10^4, 4.00 \times 10^6)$ | $1.91 \times 10^6$ ($1.16 \times 10^6, 2.86 \times 10^6$) | 1.20 (0.41, 2.06) |
| $\mu_0$ [h] | | (1, 48) | 5.55 (2.46, 9.5) | 0.44 (0.01, 1.11) |
| $\sigma_0$ [h] | | (4, 14) | 10.13 (8.43, 11.72) | 0.44 (0.01, 1.03) |
| PMF [1/48h] |  | (5, 50) | 10.56 (6.92, 17.6) | 1.75 (0.94, 2.72) |
| $E_{max}$ | OZ439 | (0.33, 0.41) | 0.35 (0.34, 0.38) | 1.08 (0.04, 2.38) |
|  | DSM265 | (0.49, 0.8) | 0.63 (0.53, 0.75) | 1.18 (0.05, 2.51) |
| $EC_{50}$ [ng/mL] | OZ439 | (16.25, 28.01) | 23.62 (19.02, 27.01) | 1.07 (0.04, 2.30) |
|  | DSM265 | (989.1, 2085.46) | 1354.4 (1088.94, 1741.38) | 1.91 (0.69, 3.15) |
| $\gamma$ | OZ439 | (1.16, 2.84) | 1.32 (1.19, 1.62) | 0.97 (0.03, 2.21) |
|  | DSM265 | (1.67, 7.27) | 2.13 (1.77, 3) | 1.35 (0.07, 2.70) |
| $k_{e0}$ [1/h] | OZ439 | (0.05, 0.17) | 0.07 (0.05, 0.11) | 1.13 (0.03, 2.52) |
|  | DSM265 | (1.7, 8.73) | 5.11 (2.5, 7.84) | 1.01 (0.03, 2.30) |
| $\alpha$ | OZ439-DSM265 | (0.2, 5) | 2.25 (0.9, 4.06) | 1.54 (0.09, 2.78) |

\* Description of the parameters and details of the prior bounds are summarised in Table 4.
<sup>†</sup> Estimated posterior samples of mono-therapy models were used to set the prior boundaries of $E_{max}$ , $EC_{50}$ , $\gamma$ and $k_{e0}$ .
<sup>‡</sup> The median and 95% credible intervals are drawn from 8,000 posterior samples (after burn-in) generated by the HMC algorithm; Supplementary Material includes the convergence metrics of HMC chains.
<sup>§</sup> Between Subject Variability (BSV) is the standard deviation of individual PD parameters from the population mean PD parameters on the logistic transform scale (see *Model fitting and simulation* section).

The results in Table 3 show that the posterior mean of the initial parasite load (sum of circulating and sequestered parasites) at the time of first parasitaemia measurement (on average 49.3 hours before administration of OZ439-DSM265), *N*_0_, was 1.91 *×* 10^6^(95% Credible Interval (CrI): 1.16 *×* 10^6^, 2.85 *×* 10^6^). The estimated initial age-distribution indicate that the majority of the parasites at the time of the first measurement were at the ring blood stage of the parasite *∼*48 h lifecycle (the posterior mean of *µ*_0_ is 5.55 hours (95% CrI: 2.46, 9.5)) with a spread (*σ*_0_) of 10.13 hours (95% CrI: 8.43, 11.72). The initial decline in parasitaemia of many of the volunteers confirms these results, since ageing of the cluster of ring parasites (observed in the blood) into trophozoites/schizonts (not observed in the blood) would lead to a temporary decline in the observed parasitaemia.

Parasite multiplication factor (PMF; the number of newly infected red blood cells by merozoites released from a ruptured shizont) was estimated to be 10.56 (95% CrI: 6.92, 17.6). *E*_*max*_ of DSM265 and OZ439 were estimated to be (0.63 (95% CrI: 0.53, 0.75)) and (0.35 (95% CrI: 0.34, 0.38)), respectively. Note that OZ439 is assumed to kill parasites at all stages [22] while DSM265 only kills trophozpoites [16], and the stage-specific action of these drugs was incorporated into the PD model (see Section *Pharmacodynamics*). Therefore, the lower estimated value of *E*_*max*_ for OZ439 compared with DSM265 does not mean that it is less potent in reducing the parasitaemia. In fact, the average of *E*_*max*_ over all the ages of the parasite’s lifecycle is 0.18 which is higher than that of DSM265 (0.14); the killing windows of DSM265 and OZ439 span 11 and 39 hours, respectively, of the 48 hours lifecycle, and the killing effect of OZ439 was halved for parasites aged 6–25 and 37–44 hours (equations (5) and (6)).

The estimated *EC*_50_ of OZ439 (23.62 (95% CrI: 19.02, 27.01)) [ng/mL] was significantly lower than that of DSM265 (1354.4 (95% CrI: 1088.94, 1741.38)) [ng/mL]. The average area under the curve (AUC) above *EC*_50_ for OZ439 were 96.4 and 103 [ng day/mL] for subjects of Cohorts A and B (OZ439 dose: 200 mg), respectively, and for DSM265 were 210.9 and 5.3 [ng day/mL] for Cohorts A (DSM265 dose: 100 mg) and B (DSM265 dose: 50 mg), respectively. The posterior median of the rate of transition between the blood plasma compartment to the effect site (hypothesised) compartment, *k*_*e*0_, was 0.07 (95% CrI: 0.05, 0.11) [1/h] for OZ439. For DSM265, the posterior median of *k*_*e*0_ was 5.11 (95% CrI: 2.5, 7.84) [1/h]. However, the distributions of prior and posterior samples for this parameter were fairly similar (Figure S5 of Supplementary Material), implying that these data were not informative for estimating *k*_*e*0_ of DSM265.

The estimated values of the OZ439-DSM265 PD interaction parameter, *α*, shows a trend toward antagonistic interaction (2.25 (95% CrI: 0.9, 4.06)). The General Pharmacodynamic Interaction (GPDI) model [23] similarly indicated an slight antagonistic interaction (*α* = 1.38 (95% CrI: 0.95, 1.95)), when the drug-drug interaction was incorporated by altering *E*_*max*_ (equation (8a)); however, when drug-drug interaction was incorporated by varying *EC*_50_ (equation (8b)), the estimated interaction was not significantly different from zero-interaction, i.e. *α* = 1.09 (95% CrI: 0.69, 1.69).

### Prediction of the optimal dosing regimen

To determine a dosing regimen that provides the WHO recommended 42-day cure rate of at least 90%, the model was simulated using the posterior samples of the individual parameters for different combinations of single doses of OZ439 and DSM265; see *Materials and Methods* section. For the parasitaemia at the time of treatment (*t* = 0), the actual values observed in malaria-endemic regions were used: the recorded parasitaemias of 1,241 patients (age range: 6 months–65 years) across 15 sites in 10 countries (Africa and South-East Asia) had a median of 52,250 (range: 2560–605,329) parasites/mL [3]. Note, only parasitaemia after drug administration is simulated, i.e. the parasitaemia growth phase was not simulated.

Figure 3 shows the mean 42-day cure rates over 20 datasets each comprised of 100 simulated hypothetical patients (2000 patients in total) who received different combinations of OZ439 and DSM265 doses; the patients whose parasitaemia got below the Lower Limit of Quantification (LLOQ: 10 [parasites/mL]) over 42 days of follow-up were considered cured. The lower and upper limits of the 42-day cure rates are shown in Figure S6 of the Supplementary Material. The dose combinations that yielded 42-day cure rates above 90% are outlined with black. The selection of the simulated doses shown in the figure fall within current evidence for safe and tolerable dosing of both drugs. OZ439 is shown to be safe and well tolerated up to 1200 mg administered as a capsule and up to 1600 mg when administered as an oral dispersion [17]. The safety profile of DSM265 is seemingly not as good as OZ439, as the number of adverse effects was higher in infected volunteers who received DSM265 compared to those who received placebo [15], although, the number of adverse events were not correlated with the administered dose.

**Figure 3:**
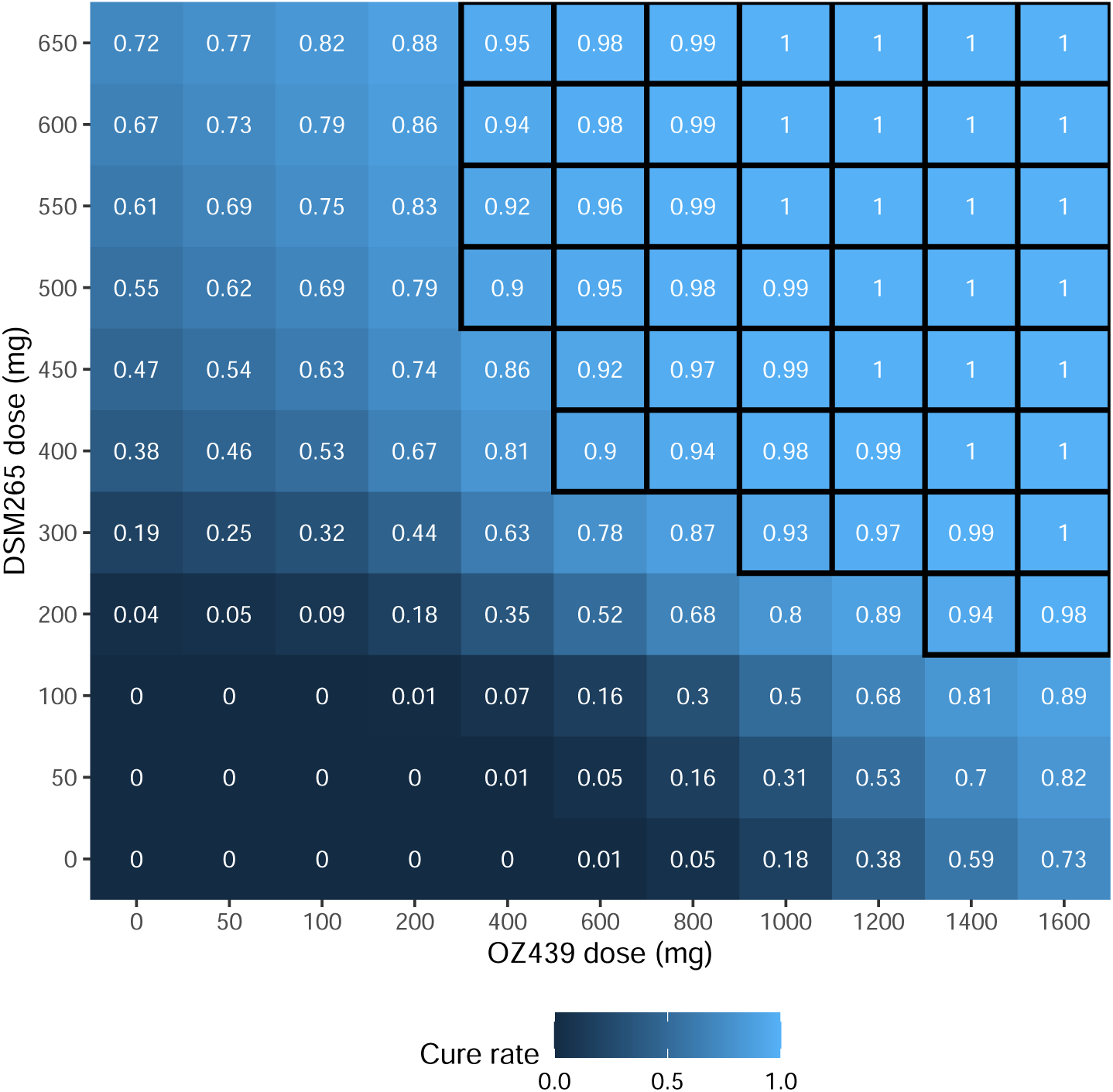
Expected cure rate within 42 days for different doses of OZ439 and DSM265 combination therapy. For each dose combination, simulations for 20 datasets each including 100 (new/hypothetical) patients were generated. The simulations were performed using the individual posterior predictive distributions of the PD parameters (see Table 3 and Section *Model fitting and simulation*) and the obtained 42-day cure rates averaged over the 20 datasets are shown in the grid squares (the lower and upper limits of the 42-day cure rates are shown in Figure S6 of the Supplementary Material). The values of parasitaemia at time of drug administration in the simulations were drawn from a log-normal distribution constructed using the reported values in malaria-endemic countries: median = 52,250 (range: 2560–605,329) parasites/mL [3]. Single doses of OZ3439 and DSM265 were administered at times 0 and 2 h, respectively. The dose combinations that yielded a 42-day cure rate ≥ 90% are outlined with black borders. Cure rate: proportion of patients in each dataset that had parasitaemia below the LLOQ (10 [parasites/mL]) over 42 days of follow-up.

The results in Figure 3 show that the 42-day 90% cure rate cannot be achieved for doses up to 200 mg of OZ439 and 100mg of DSM265. Various dose combinations achieve a ≥ 90% 42-day cure rate, however, considering safety and tolerability of each drug, we recommend the combination of 800 mg of OZ439 and 450 mg of DSM265 be investigated in further trials. This dosing regimen also provides high drug concentrations of OZ439 in the blood plasma, covering the exposure duration of DSM265 for up to 42 days.

The initial distribution of the age of parasites for each simulated patient was assumed to follow that estimated from the volunteers. A sensitivity analysis, where the mean of the initial parasite age distribution was assumed to follow a uniform distribution over (0, 48 h), was performed (see Figure S7 of the Supplementary Material). The results showed that predicted cure rates can vary slightly by changing the initial parasite age distribution, however, for 800 mg of OZ439 and 450 mg of DSM265 the predicted 42-day cure rate remained above 90%, confirming the robustness of this dosing regimen.

## Discussion

We proposed a within-host mathematical PK-PD model for the combined antimalarial activity of two novel drugs: OZ439 and DSM265. Our model incorporated the parasite age-specificity of killing action of the drugs, parasite sequestration and PD interaction between the drugs. Using a Bayesian hierarchical framework, our model provided a good fit to parasitaemia data collected pre- and post-administration of OZ439 and DSM265 from healthy volunteers innoculated with *P. falciparum* malaria. Simulating parasitological outcomes using the estimated PD parameters determined the safe and tolerable dosing regimens of 800 mg for OZ439 and 450 mg for DSM265 can yield the WHO recommended 42-day cure rates (*≥* 90%).

Our model simulations put forward a set of potential OZ439-DSM265 dose combinations that are predicted to be efficacious and are within the safety and tolerability limits [15, 17]. Among these dose combinations, one must be selected for deployment that does not lead to development of parasite resistance to one of the drugs due to long durations of sub-therapeutic exposure. The significantly longer exposure time of DSM265 (see Figure 1) reinforces the possibility of resistance development to this drug, if the parasites are not simultaneously exposed to another drug. In fact, a study by Llanos-Cuentas et al. 2018 [24] found evidence of selection of resistance to DSM265 through a mutation in the DHODH enzyme target in Peruvian patients who were administered a single dose of DSM265. Therefore, the selected dose of OZ439 must be sufficiently high such that it exposes the parasites for long enough timespans during which the parasites are exposed to DSM265 as well. As a result, the selection of higher doses of OZ439 must be favoured from the set of all efficacious, safe and tolerable doses (Figure 3), e.g. 600mg or 800mg of OZ439, combined with 400 mg or 450mg of DSM265.

We proposed a novel model of the combined action of the drugs based on the Bliss independence concept [25]. Fitting the model to the parasitological data showed that OZ439 and DSM265 have a slight antagonistic interaction, which was also confirmed using a GPDI model for drug interaction; however, the antagonistic interaction was not strong enough to significantly nullify their combined effect, and the combination compound was still able to produce cure rates above 90%. We showed that the predicted cure rates can be influenced by the assumption about the initial age distribution of the parasites – highlighting the significant influence that synchronicity of infection at admission can have on treatment’s efficacy and the importance of incorporating that into a mathematical model – however, the suggested dosing regimen (800mg of OZ439 combined with 450mg of DSM265) still provided a *≥* 90% cure rate.

The PK-PD model proposed in this work can be used to guide phase 2 and 3 clinical trials evaluating the efficacy of OZ430–DSM265 regimens, helping to reduce the financial and logistical costs of these trials. The model did not include the potential contribution of host immunity to parasite clearance [26] since it was validated on volunteers who had not been previously exposed to malaria infection. However, the influence of immunity on parasite clearance would augment drug effect therefore resulting in an overestimate of the minimal efficacious dose, and thereby result in a greater safety margin. The efficacy of the suggested dosing regimen in reducing gametocytaemia, and thereby transmission, was not investigated in this work. This will be considered in future work based on a model we have developed for within-host transmission dynamics [27]. Further, a more mechanistic model of the combined action of the drugs that accommodates the underlying processes of drug interaction could be used [28]. However, to do so would require a greater understanding of the drug-drug interactions that is yet unavailable, requiring more sophisticated *in vitro* parasite susceptibility experiments, e.g. checkerboard assays, that focus particularly on the combined effect of the drugs.

*P. falciparum* parasites resistant to ACTs are rapidly spreading across South-East Asia, impeding the goal of WHO to achieve malaria elimination by 2030 in this region. The combination of OZ439 and DSM265, administered according to the suggested efficacious and well tolerated regimens, appears to be a promising alternative treatment to replace the failing ACTs.

## Materials and Methods

### Data

Data from four separate studies of volunteers innoculated with *P. falciparum* malaria were used for estimating the parameters in this work: (i) OZ439 mono-therapy (doses: 100, 200 and 500 mg) [12]; DSM265 mono-therapy (doses: (ii) 150 mg [14] and (iii) 400 mg [15]);(iv) OZ439-DSM265 combination therapy (doses: 200 mg of OZ439 combined with 50 and 100 mg of DSM265) [7]; details of these studies are summarised in Table 1. The data from the mono-therapy studies were used for constructing the prior distributions of the PD parameters, as detailed in the *Model fitting and simulation* section.

In the studies of OZ439 mono-therapy and OZ439-DSM265 combination therapy, the volunteers were initially inoculated with *∼*1800 *P. falciparum-*infected red blood cells and were admitted and confined for 48 hours before the compounds were administered, after their parasitaemia reached *≥* 1000 parasites/mL or clinical symptoms appeared (whichever occurred first). In the DSM265 mono-therapy where 150 mg dose was administered, the volunteers were inoculated with *∼*1800 and the threshold for admission was considered to be 800 parasites/mL [15]. In the other DSM265 mono-therapy where 400mg of DSM265 was administered, the volunteers were inoculated with *∼* 2800 viable *P. falciparum* parasites and were treated on day 7 [14]. A single dose of the compounds were administered in all of the studies. All the volunteers received rescue treatment on a certain day following the drug administration or after parasitaemia recrudescence; only the parasitaemia measurements before the rescue treatment was administered were included in the analyses.

### Mathematical model

The within-host PD model fitted to the parasitaemia data was based on the models of [20, 21], which include the stage-specificity of drug action, shown in susceptibility experiments to significantly impact killing effect of antimalarial drugs [29, 30]. Interaction between the PD action of the drugs was also incorporated in the model to capture the combined effect of OZ439 and DSM265.

### Pharmacokinetics

A two-compartment PK model with first-order absorption for OZ439 and zero-order absorption for DSM265 best described the PK profiles of the volunteers. The pharmacokinetics of each drug were not altered when the drugs were given in combination [7]. A delayed effect of the plasma drug concentration of both OZ439 and DSM265 on parasite killing was incorporated in the model. This was modelled as a transition between two compartments with rate *k*_*e*0_; see Supplementary Material for further information about the PK models. The concentrations at effect site, *C*_*e*_(*t*), were substituted into the PD model to derive drug action, as detailed below.

### Pharmacodynamics

The PD model in [6] was used for the time-evolution of the number of parasites in the body, *N*:

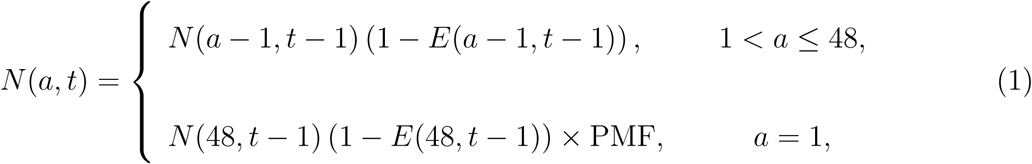

where *a* is the parasite age, taking only integer values over the range 1 to 48, *t* is time, taking only integer values, and PMF is the *parasite multiplication factor*, which represents the number of merozoites released into the blood by a shizont at the end of its lifecycle which successfully invade red blood cells. *E*(*a, t*) is the killing effect of the drug, taking values between 0 and 1, and dependent upon the age of parasites during [*t, t* + 1). The subjects were assumed to be infected with an initial parasite load of 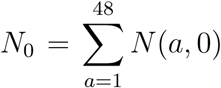 which has a discretised normal distribution over age with the mean at *µ*_0_ hours and standard deviation *σ*_0_ hours (both on the continuous scale) and *N* (1, 0) = PMF*×N* (48, 0). To determine *N* (*a*, 0) by discretising a continuous normal distribution, *n*(*a*) *∼ 𝒩* (*µ*_0_, *σ*_0_), the following formula was used

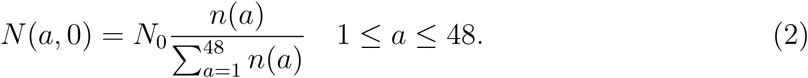

The number of detectable parasites circulating in the blood, *M* (*t*), is determined by

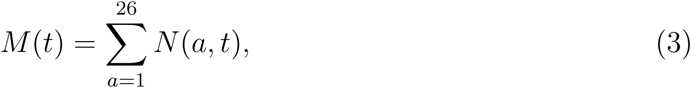

where it is assumed that the infected red blood cells in circulation (the parasitaemia measured by blood samples) constitute mostly of ring stage parasites, because the older parasites sequester in blood capillaries and are not visible in the blood samples [31]. The number of parasites per mL of blood (the unit of parasitaemia in the data) was determined by dividing *M* (*t*) by each patient’s blood volume in *mL*; the patients were assumed to have 70 *mL/kg* blood, hence patient’s blood volume was calculated as 70 *mL/kg ×* patient’s weight.

We assumed Michaelis-Menten kinetics for *E*:

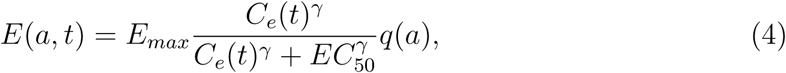

where *E*_*max*_ is the maximum killing effect of DHA; *C*_*e*_(*t*) is the drug concentration at the effect site at time *t*; *EC*_50_ is the concentration at which 50% of the maximum killing effect is obtained; *γ* is the sigmoidicity (also known as slope) of the concentration-effect curve. The stage specificity of the killing effect is applied using the *q*(*a*). A description of all the PD parameters are provided in Table 4.

**Table 4:** Description and selected prior bounds (mono-therapies) for each parameter of the PD model.

| Parameter | Trial | Prior bounds | Description |
| --- | --- | --- | --- |
| $N_0$ | OZ439<br>DSM265 | $(5.12 \times 10^4, 8.02 \times 10^6)$<br>$(5.18 \times 10^4, 1.42 \times 10^7)$ | initial number of parasites (circulating + sequestered)<br>*lower bound*: $\text{LLOQ} \times \text{mean blood volume of volunteers in the trial}$<br>*upper bound*: $2 \times \text{maximum observed parasitaemia at the first blood measurement} \times \text{mean blood volume of volunteers in the trial}^*$ |
| $\mu_0$ | OZ439<br>DSM265 | (1, 48) | mean of initial parasites age distribution [h]<br>*bounds*: lowest and highest ages of parasites in a 48h lifecycle |
| $\sigma_0$ | OZ439<br>DSM265 | (4, 14) | standard deviation of initial age distribution [h]<br>*bounds*: selected to produce a wide range of dispersion (narrow to dispersed) in initial age distribution |
| PMF | OZ439<br>DSM265 | (5, 50) | parasite multiplication factor [/48 h]<br>*bounds*: according to [40] |
| $E_{max}$ | OZ439<br>DSM265 | (0.05, 1) | maximum killing effect<br>*bounds*: an arbitrarily wide range is selected |
| $EC_{50}$ | OZ439<br>DSM265 | (0.5, 50)<br>(500, 5000) | concentration producing $E_{max}/2$ effect [ng/mL]<br>*bounds*: according to [12] and [16] for OZ439 and DSM265, respectively |
| $\gamma$ | OZ439<br>DSM265 | (1, 10) | sigmoidicity of the concentration-effect curves<br>*bounds*: selected to generate a wide range of sigmoidicities |
| $k_{e0}$ | OZ439<br>DSM265 | (0.01, 10) | rate of drug transition from blood [1/h]<br>plasma compartment to the effect-site compartment<br>*bounds*: arbitrarily large range was selected |
| $\alpha$ | OZ439-DSM265 | (0.2, 5) | interaction parameter<br>*bounds*: the selected values produce a wide range of interactions, from strong antagonism to strong synergism |

\* see Section *Model fitting and simulation* for conversion of the observable circulating parasitaemia (no. parasites/mL) to the total parasite biomass (circulating + sequestered)

Previous *in vitro* experiments showed that OZ439 kills the parasites at all the stages of the blood lifecycle [13], and DSM265 only kills trophozoites [16]. It has also been shown that OZ439 has its maximum activity against trophozoites (reviewed in [22]). Therefore we considered the following step functions for the stage specificity of the killing action of the drugs:

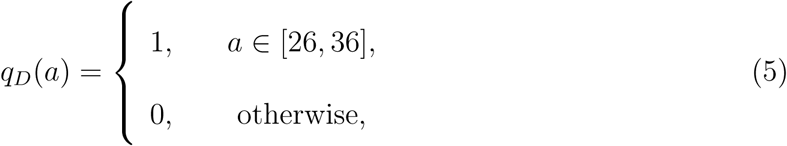

and

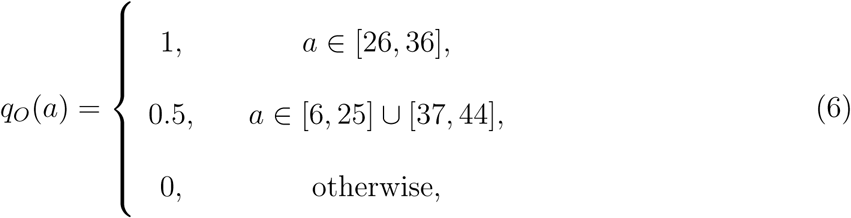

where *q*_*D*_ and *q*_*O*_ are the stage-specificity functions for DSM265 and OZ439, respectively. Note, early ring and late schizont parasites were considered insensitive to OZ439, similar to artemisinin [32].

### Combined killing effect

An empirical approach was taken to model the combined effect of OZ439 and DSM265. Considering the unavailability of *in vitro* parasite susceptibility data, the number of interaction parameters were kept to a minimum. In addition, we selected a model where the combined effect monotonically increases with concentrations of OZ439 and DSM265, since, except in very rare cases, increasing the concentration of each drug must either increase the combined effect or the combined effect remains unchanged.

To characterise the interaction between OZ439 and DSM265 and model their combined action, a zero-interaction framework must first be defined. Two widely used empirical frameworks for zero-interaction are *Loewe additivity* [33] and *Bliss independence* [25]; the former is used when the drugs are believed to have similar modes of action and the latter is when the drugs act through completely different mechanisms; see [6, 34] for further information. Different types of drug-drug interaction (synergism/antagonism) can then be modelled by characterising deviation from the zero-interaction model and the combined killing action can be defined accordingly.

OZ439 and DSM265 have different modes of action – the former kills the parasites by activating the endoproxide bond [12] and the latter by inhibiting the parasite’s DHODH enzyme [15] – hence Bliss independence was selected as the base model for zero-interaction. The Bliss independence model was then modified to define the combine effect, *E*_*OD*_:

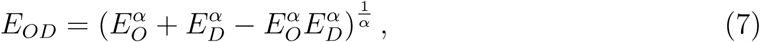

where *E*_*O*_ and *E*_*D*_ can be obtained using the Michaelis-Menten function (equation (4)), and *α* is the interaction parameter. The values of *α* = 1, *α >* 1 and 0 *< α <* 1 correspond to zero-interaction, antagonism and synergism, respectively. The combined effect defined in this form is monotonically increasing and has a well-behaved form in regard to *α*. Figure 4 depicts the combined effect for three different types of drug-drug interaction.

**Figure 4:**
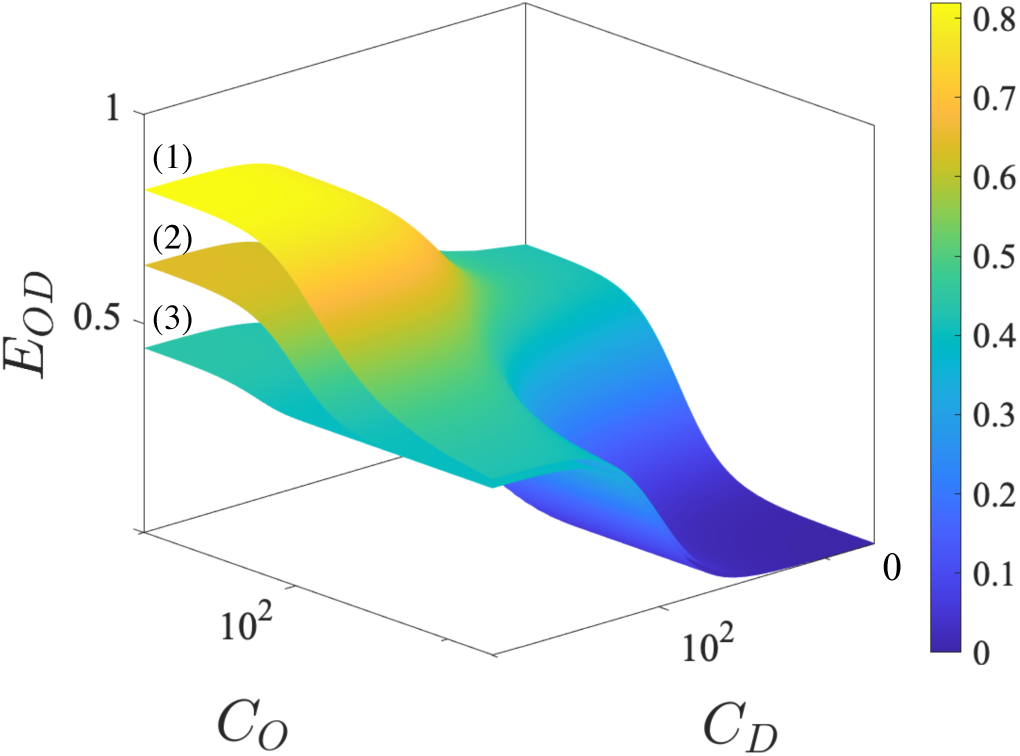
Combined effect of OZ439 and DSM265, *E*_*OD*_ in equation (7), for three different types of interaction and the following arbitrarily selected set of PD parameter values: *E*_*max,D*_ = *E*_*max,O*_ = 0.4; *γ*_*D*_ = 3; *γ*_*O*_ = 3; *EC*_50,*D*_ = *EC*_50,*O*_ = 100 [*ng/mL*], where *O* and *D* in the sub-indices denote correspondence to OZ439 and DSM265, respectively. The (1), (2) and (3) surfaces correspond to *α* = 0.3 (synergism), *α* = 1 (zero-interaction) and *α* = 7 (antagonism).

In addition to the previous approach for accommodating interaction (equation (7)), a simplified version of the GPDI model proposed by Wicha et al. [23] was examined. In their proposed model, *EC*_50_ and *E*_*max*_ are scaled depending on the concentration of the other drug and nature of the interaction (i.e. synergism/antagonism). However, the full form of their model was not used here due to potential non-identifiability of the parameters as well as non-monotonicity of the combined effects. To implement the simplified version of the GPDI model, *E*_*max*_ and *EC*_50_ are modulated as below

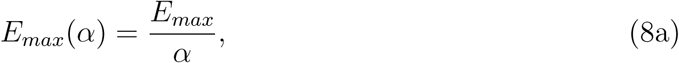

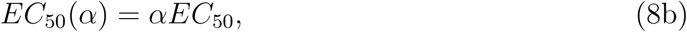

where *α* is the interaction parameter, and *α* = 1, *α >* 1 and 0 *< α <* 1 correspond to zero-interaction, antagonism and synergism, respectively. *E*_*max*_(*α*) was bounded to be ≤ 1 (see equation (1)).

### Model fitting and simulation

A sequential approach was employed to fit the model to data: the PK parameters were first estimated to simulate drug concentration profiles which were then substituted into the PD model to estimate the PD parameters. Fitting the PK models to the data was performed in a non-linear mixed effects modelling framework using Monolix [35] as follows. The individual PK parameters were estimated using the mode of the conditional distribution of the individual parameters. A total of 500 exploratory and 200 smoothing samples were generated using the Stochastic Approximation Expectation-Maximization (SAEM) method to estimate the population PK parameters. Finally, linearisation method was used to estimate the Fisher Information Matrix and the log-likelihood.

The PD model was fitted in a Bayesian hierarchical framework that allowed estimating individual parameters and incorporating the prior information about the parameters. The likelihood function was formed by assuming that the log-transformed parasitemia of the subjects have a normal distribution with mean at the simulated parasitaemia and a certain standard deviation. For parasitaemia below the quantification level, the M3 method was used, by considering the data below the quantification level left-censored and using the cumulative normal distribution in the likelihood function [36]; see Supplementary Material for further information.

The joint posterior distribution of the parameters was sampled using the Hamiltonian Monte Carlo (HMC) method [37]. Four chains were initialised randomly from different points to sample the posterior distribution. A total of 4,000 samples were generated by each chain, half of which were discarded as warm-up, leaving 8,000 samples in total from which to draw our inferences. The RStan package [38] in the R software [39] was used to implement the HMC method.

The individual parameters were logistic transformed from their original bounded do-main to a new unbounded domain, using

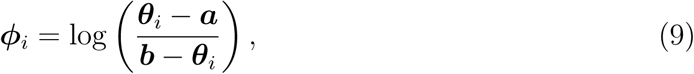

where ***θ***_*i*_ represents an individual model parameter in the original domain, and ***a*** and ***b*** are the lower and upper bounds of ***θ***_*i*_. A multivariate normal distribution with mean ***ϕ*** and covariance matrix **Ω** was considered for ***ϕ***_*i*_. The generated posterior samples of ***ϕ***_*i*_ were mapped back using the inverse function of equation (9) to get ***θ***_*i*_ over (***a, b***). The interaction parameter, *α*, was log-transformed in order to generate the same proportion of prior samples of *α* producing antagonistic and synergistic interactions, and then back-transformed to be used in equation (7); see the Supplementary Material for further information.

The values of *a* and *b* for the combination therapy and mono-therapies are listed in Tables 3 and 4, respectively. In fitting the model to data of mono-therapy trials, wide ranges that contain all feasible values for parameters were used. For fitting the model of combination therapy to the data, the results of the mono-therapy models were used to set the prior bounds of *E*_*max*_, *γ, EC*_50_ and *k*_*e*0_. To be specific, the 2.5% and 97.5% percentiles of the obtained posterior parameter samples in the mono-therapy models were used as *a* and *b*, respectively. This was considered assuming that the values of these parameters mostly depend on the activity of the individual drugs, hence their values in the combination therapy should be very close to those in the mono-therapies.

Convergence of the HMC chains were assessed by evaluating the following metrics: i) the potential scale reduction statistic,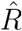, which shows how well the chains are mixed – satisfactory convergence of chains yields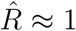; ii) effective sample size, *n*_*eff*_, is an estimate of the number of independent draws, after accounting for autocorrelation between the posterior samples.

Model simulations for predicting the cure rates were performed using the posterior samples of individual parameters. The cure rate was defined as the proportion of 1000 patients whose parasitaemias were below the LLOQ (10 [parasites/mL]) within 42 days were considered cured. For the individual PD parameters, first, 20 samples of ***ϕ*** and **Ω** were selected from the set of 8000 posterior samples – the last five posterior samples from each of the four HMC chains were used. Using the selected samples, 100 samples of ***ϕ***_*i*_ were generated from a multivariate normal distribution with mean ***ϕ*** and covariance matrix **Ω**. The generated samples of ***ϕ***_*i*_ were then back-transformed using the inverse function of equation (9) to get the original PD parameter samples, ***θ***_*i*_, which were used to simulate 20 cohorts/datasets each including 100 hypothetical patients. (see Section 4 of Supplementary Material). Note, the total number of parasites was set to zero if it reached values below 1 in the simulations. For simulation of the PK profiles of OZ439 and DSM265, samples were generated using the distributions, population parameters and between subject variabilities listed in Table 2.

The parasitaemias observed in the field were used as the baseline parasitaemia in the simulations; according to [3], the distribution of parasitaemias of 1,241 patients across 15 sites in 10 countries had median of 52,250 parasites/mL and spanned the range of 2560–605,329 parasites/mL. Thus, a log-normal distribution with the geometric mean at 52,250 parasites/mL and standard deviation on the log-scale of 0.78 = (log(605329) *−* log(2560)) */*7 were used for generating samples of baseline parasitaemia in the simulations. The total circulating parasitaemia, *M* (0), was obtained by multiplying the generated samples from the log-normal distribution to average blood volume of the volunteers in the combination therapy trial, 5036.77 mL. To obtain the total parasite burden for an individual, i.e. sum of the sequestered and circulating parasites (*N*_0_), we used the posterior samples of *µ*_0_ and *δ*_0_ estimated for the volunteers as the mean and disperse of the initial age distributions. Finally, the age distribution and *M* (0) were substituted into equations (2) and (3) to obtain *N*_0_ for each simulated subject.

## Acknowledgements

This work is supported in part by the Australian Centre for Research Excellence in Malaria Elimination, funded by the NHMRC (1134989). JAS is funded by an Australian National Health and Medical Research Council of Australia (NHMRC) Senior Research Fellowship (1104975). JSM is funded by a NHMRC Program Grant (1132975) and Practitioner Fellowship (1041802). The clinical trials (NCT02389348, NCT02573857, ACTRN12613000522718, ACTRN12613000527763 and ACTRN12612000814875) from which the data were derived were supported by the Medicines for Malaria Venture (MMV) and funded by the Wellcome Trust (grant reference number: 095909/Z/11/Z), a grant by the Global Health Innovation and Technology Fund (GHIT) (grant no. G2014-108), and by funding from the Bill and Melinda Gates Foundation.

NG, MC and JJM are employed by MMV; none of the other authors declares any competing interests.

